# Towards a unification of niche and neutral models of community ecology

**DOI:** 10.1101/273664

**Authors:** Andres Laan, Gonzalo G. de Polavieja

**Affiliations:** Champalimaud Research, Champalimaud Center for the Unknown - Lisbon, Portugal

## Abstract

Ecological models of community dynamics fall into two main categories. The neutral theory of biodiversity correctly predicts various large-scale ecosystem characteristics such as the species abundance distributions. On a smaller scale, the niche theory of species competition explains population dynamics and interactions between two to a dozen species. Despite the successes of the two theories, they rely on two contradictory assumptions. In the neutral theory each species is competitively equivalent while in the niche theory every species is specialized to exploit a specific part of its environment. Here we propose a resolution to this contradiction using a game theory model of competition with an attractor hyperplane as its equilibrium solution. When the population dynamics shifts within the hyperplane, it is selectively neutral. However, any movement perpendicular to the hyperplane is subject to restoring forces similar to what is predicted by the niche theory. We show that this model correctly reproduces empirical species abundance distributions and is also compatible with species removal experiments.

## Introduction

Current theoretical descriptions of community ecology and biodiversity are split between two different approaches. Large scale processes are typically analyzed using the unified neutral theory of biodiversity [1, 2]. Neutral theory has successfully explained complex statistical patterns of species abundance distributions observed in ecological communities such as tropical forests [3] and river deltas [4]. Despite its success in explaining large-scale patterns, it relies on assumptions which do not hold at the species scale.

According to neutral theory, all species are competitively equal and differ only in abundance. Field observations find this assumption repeatedly violated. Species occupy preferred niches [5], they have differential effects on community dynamics [6, 7] and exhibit reliable recovery from perturbations [8]. Such results are better explained by models where each species has a competitive advantage over others within a distinct niche [9]. Mathematically, the interactions between species are then described by non-linear population dynamics models rather than stochastic drift equations [1, 5, 9].

This divergence between the two theories in underlying assumptions and mathematical apparatus is obviously problematic and there have been several calls to unify the two descriptions [10]. Previous steps in that direction include theories of emergent neutrality [11] and models which introduce partial competition into neutral theories [12]. Despite these advances, uncertainty remains about the degree to which the attempts have been successful [10].

Here, we propose a game theory model of large-scale ecosystem dynamics which is compatible with several key empirical findings on community dynamics. Our model has as its equilibrium solution an attractor hyperplane. Within the hyperplane, dynamics are neutral and the model is thus compatible with stochastic drift-based ecological dynamics models. We find that just like the neutral theory, our model successfully reproduces species abundance distributions of Panamanian trees [13] and Mediterranean phytoplankton [14]. Under a particular parametrization, it is also consistent with the finding that species distributions may occasionally be weakly multi modal [15]. When our model is perturbed with stimuli that move the system off the hyperplane, dynamics are no longer neutral. Instead, the perturbation is resisted by competitive processes. The restorative dynamics of the model are compatible with species removal experiments [8, 16]. Our model may thus represent a promising mathematical framework which can unify the key empirical findings of ecosystem dynamics without internal contradictions.

## Results

In the search for a unification of the neutral and niche theories, we took inspiration from the hawk-dove game studied in evolutionary game theory [17, 18]. The hawk-dove game is characterized by two strategies: the hawk strategy and the dove strategy. Hawks and doves mix in the population randomly and compete against one another. When two hawks meet, both suffer a fitness loss of 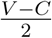(where V is the value of the resource they are competing for and C is cost of mutual competition, also *C > V >* 0). On the other hand, when a hawk meets a dove the dove surrenders and the hawk increases its fitness by V while a dove gets nothing. When two doves meet they split the resource and each gets 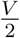 increase in fitness. The game is at equilibrium if the frequency of hawks in the population is equal to 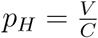.

In our extension, each species participates in *N* hawk-dove games simultaneously. Each game can be thought as being a competition for a different kind of essential resource. Game 1 might correspond to competition for light, game 2 for competition for nitrogen and so on. Each game has its corresponding parameters, for game *i* with values *V*_*i*_ and *C*_*i*_.

Each species must choose a micro strategy (either play hawk or play dove) for each of the *N* games. For the case of 4 games, for example, a macro strategy for an individual specifies how to play in the 4 games. An example macro strategy may be *H*_1_ *D*_2_ *D*_3_ *H*_4_, which specifies to play hawk in game 1, dove in game 2, dove in game 3 and hawk in game 4. Furthermore, we postulate that if two individuals have exactly the same macro strategy then they belong to the same species. The total payoff of a macro strategy is determined by its cumulative pay-off in all of the *N* games.

For *N* games there is a total of 2^*N*^ different macro strategies. The state of the population is then characterized by 2^*N*^ probability values *π*_*j*_, where *π*_*j*_ is the probability of encountering macro strategy *j*. The population is at equilibrium when every strategy has the same expected fitness. Since all *N* games are independent from one another in terms of pay-offs, then the population must be in equilibrium with respect to each of the *N* games. Therefore, the probability of encountering the hawk strategy in game *k* (*p*_*H*_*k*__) must be equal to 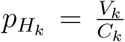. We can write that condition in terms of the probabilities of the composite strategies as 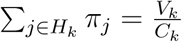, where *j∈H*_*k*_ stands for all the strategies *j* which play hawk in game *k*. We get one such equation for each of the *N* games and another equation which requires that all the probabilities sum to 1. Therefore, we end up with *N* + 1 equations for 2^*N*^ variables and the final solution is under-determined. In effect, the mathematical structure of the solution is constrained to lie in somewhere inside a 2^*N*^ *-N-* 1 dimensional hyperplane. Within that hyperplane, all solutions are equally good and therefore there is plenty of room for neutral drift between the strategies. We tested whether the model reproduces the species area distribution in the Barrio Colorado tree community (BCI)a standard large-scale dataset for community ecology [13]. Since the community contains around 200 species, we chose a value of *N* = 8 and we sampled each *V*_*i*_ independently and uniformly at random in the range from 1 to 1.9. The starting probability distributions of all 2^*N*^ species were chosen from a uniform random distribution and normalized to sum to one. We simulated the replicator equation in various random starts and observed the equilibrium state distributions of strategy probabilities frequently converging to the log-normal distribution, which is very different from our starting state- the uniform probability distribution. The same also held true if we initialized the population from a Gaussian distribution. We found an excellent match between the simulation and empirical data (Figure 1, left). We obtained a similar result for a Mediterranean phytoplankton dataset [14, 19] (Figure 1, right).

**Figure 1:**
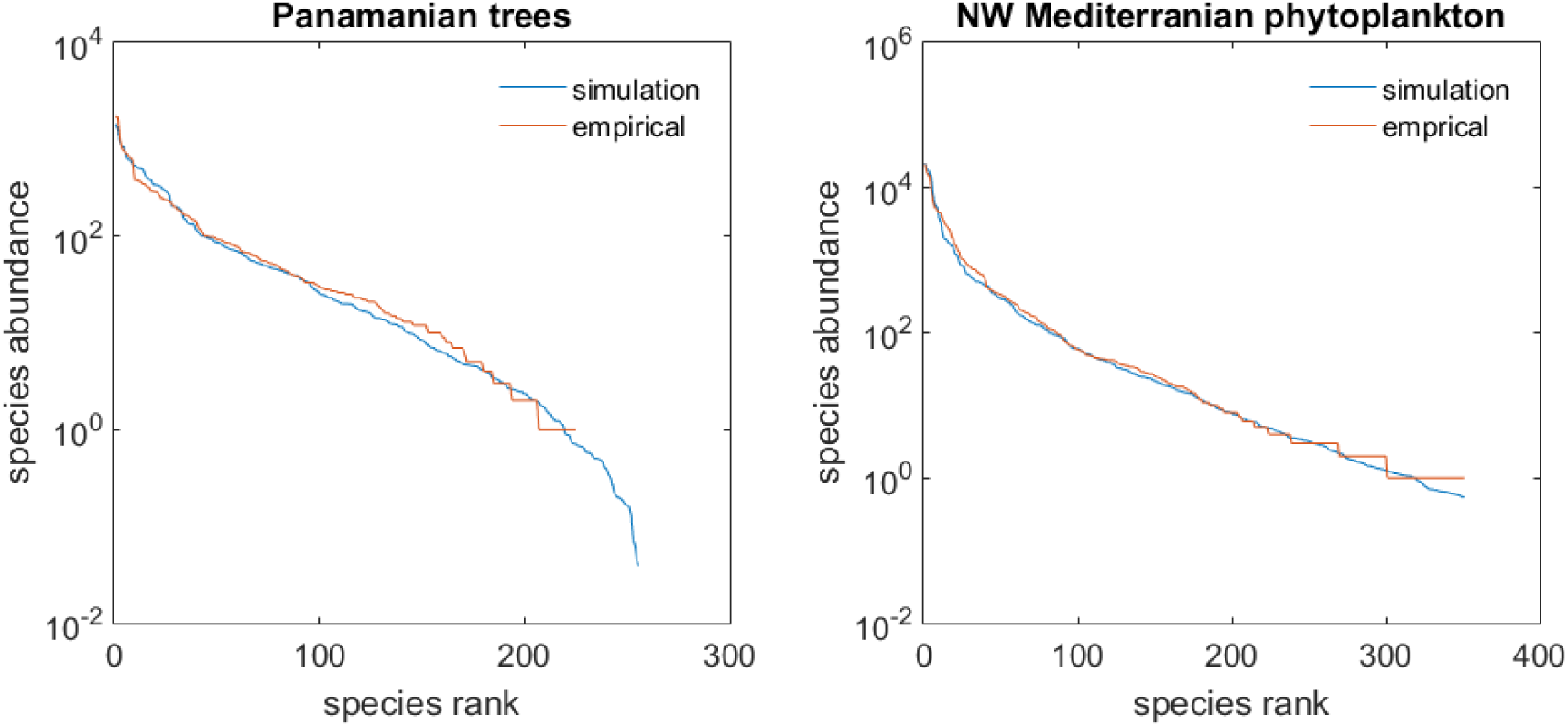
Species abundance distributions (SADs). SADs for two datasets- the Panamanian trees dataset [13] (left panel) and the North-West Mediterranean phytoplankton [14] (right panel). Red: empirical distributions. Blue: Model results at equilibrium. Simulation parameters for BCI dataset as given in main text. For the phytoplankton, we used *N* = 9 (needed because we have more species) and V was sampled uniformly at random between 1 and 1.99 to generate greater variance. Furthermore, a cut-off at 350 species was used when presenting the histogram to better facilitate comparison with data.

A mathematical analysis of our model shows that how the log-normal distribution emerges as a solution (**Appendix**). Some species abundance distributions have been described as having some degree of multi-modality [15]. In this respect, it is intriguing to note that under certain distributions of *V*_*i*_ values, multi-modal species distributions emerge in our model as well (**Appendix**).

Previously, several other mathematical models have shown a good match with the BCI dataset. One model has been the so-called null model-the log-normal distribution [20]. It provides excellent fits, but it does not specify a theory of how it arises [21]. Another model is the zero-sum multinomial distribution. It emerges as a solution to the neutral theory of biodiversity [1]. However, this theory is problematic not because it does not fit species abundance distributions, but because it assumes that all species are competitively neutral. Several experiments have demonstrated that this is not the case [5, 8, 10].

In particular, we considered species removal experiments [8, 16, 22]. In these experiments, some members of a particular species are removed from the community such that the prevalence of that species falls below its usual levels. Then, the reproductive success of that species is compared in the reduced conditions and the common conditions. The usual finding is that when a species level has been reduced, it experiences a temporary competitive advantage compared to other species and its prevalence tends to move back towards its original level. This finding contradicts the neutral theory of biodiversity.

Our model is compatible with species removal experiments. To see why, let us assume that we reduce the total number of one strategy type, let’s exemplify it with the *H*_1_ *D*_2_ *D*_3_ type. Then, we have created a deficiency in the population of *H*_1_, *D*_2_ and *D*_3_ strategies and the three experience restorative forces. Some members of the population may also experience an increased fitness (for example the *H*_1_ *D*_2_ *H*_3_ probably experiences a net boost in levels due to its first 2 components), but *H*_1_ *D*_2_ *D*_3_ is the only one which experiences increased fitness along all its three components and therefore has the largest fitness advantage.

We simulated this scenario (**Figure 2**). Following the experimental study from [8], we removed half of the members of a species and observed its recovery. We plot the results of 600 simulations. In each, we first let the population converge to an equilibrium, then we removed half the members of one species, and we let the population recover. We found there is always recovery with respect to the perturbation (red points are above the blue points) although the recovery is not perfect (red lines versus yellow line). Intriguingly, the more prevalent a species is initially, the better its levels recover. This result qualitatively mirrors what is observed in plant community dynamics [8]. Therefore, unlike the neutral theory of biodiversity, our theory is not directly contradicted by community perturbation experiments and may represent a way forward towards a more realistic unification of neutral and niche dynamics.

**Figure 2:**
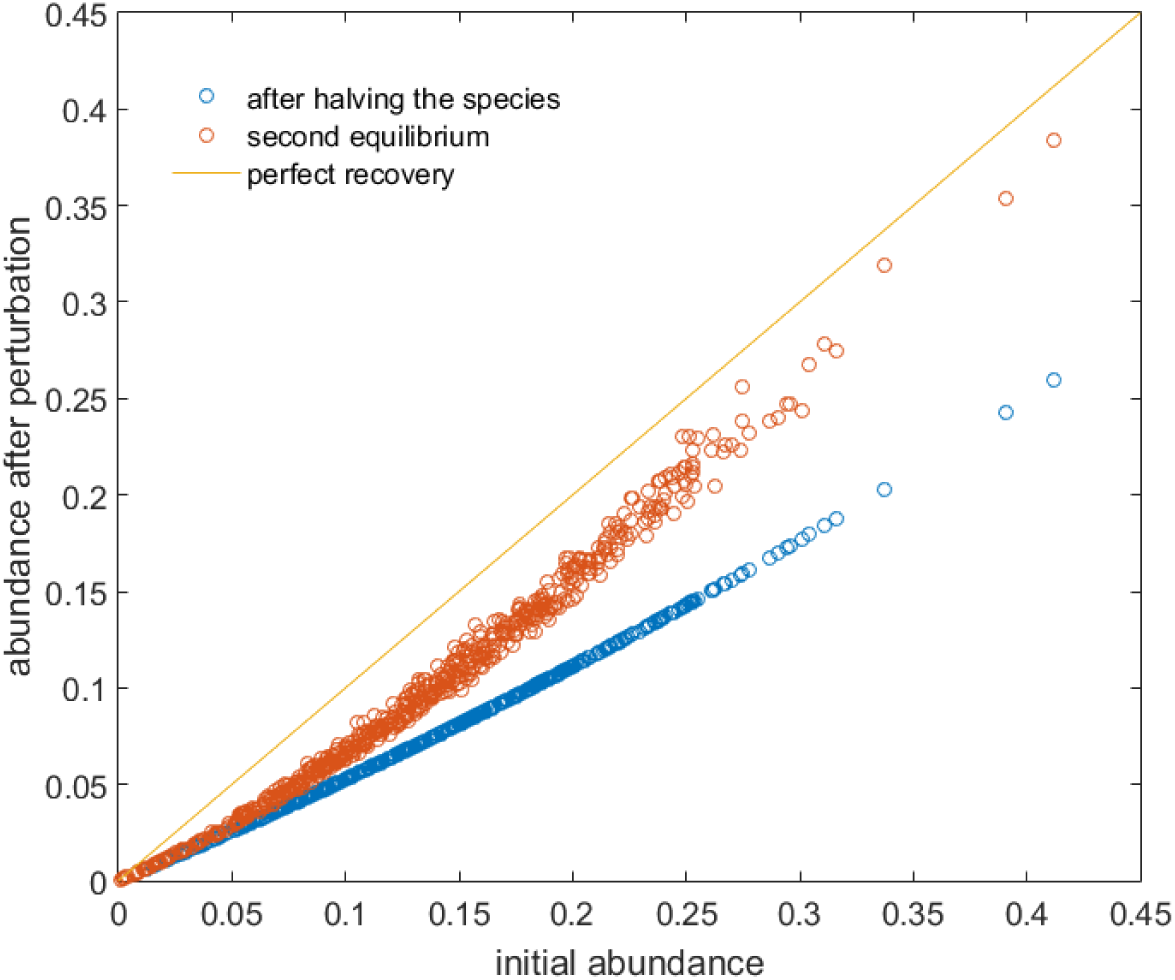
Simulation of removal experiments. The blue points plot the prevalence of a species after its numbers were halved against its initial prevalence. The red points show population recovery after a new equilibrium has been reached. Yellow line shows hypothetical perfect recovery.

## Discussion

We have proposed a model of species interactions which holds the potential to unify the niche and neutral perspectives on biodiversity [10]. Different from other attempts to combine niche and neutral theories, we do not need to model factors like species creation, immigration from a surrounding community or dispersal limitations [12, 23]. Even in a fully mixed population with no source of new species, we obtain an approximately log-normal species abundance distribution in equilibrium. Due to the attracting nature of the hyperplane, the equilibrium is also reached very fast which may be an advantage over other models, where kinetic equilibriums can be slow to emerge and are transient in nature [11]. In the future, it will be interesting to incorporate such considerations into our baseline model to facilitate further comparisons between models. The central object which enabled us to achieve this in our model was the hyperplane attractor structure of our model. We anticipate that even if the details of our model prove to be inaccurate in some respects, this underlying mathematical structure may nevertheless be present in any model which might end up replacing our current formulation. This is likely to be the case because the only way that neutral drift and restorative dynamics can simultaneously co-exist is if dynamics is neutral along some directions but restoring on other- the very property which defines an attractor manifold. Similar mathematical structures have previously proved useful in analyzing other complex networks such as neural dynamics [24]. We hope that our model will stimulate further research into attractor hyperplanes in theoretical ecology as well.

## Appendix

### Approximate analytical solution

Here, we show that an approximate solution to our model can be expressed in terms of the solution to the individual component hawk dove game models. We define the following notation. *π*_*i*_(*t*) is the probability of observing macro strategy i at time t. *p*_*H*_*k*__(*t*) is the the probability of encountering a hawk player in game *k* at time *t*. As before, *p*_*H*_*k*__(*t*) = *j*_*∈*_ *Hk π*_*j*_(*t*), where *j ∈H*_*k*_ stands for a strategy j which includes playing hawk in game k as its element. *s*_*H*_*k*__(*t*) stands for the function which solves the two-player hawk dove game replicator equation with pay-offs *V*_*k*_, *C*_*k*_ and where *s*_*H*_*k*__(0) = *p*_*H*_*k*__(0). In other words, *s*_*H*_*k*__(*t*) would capture the the time evolution of the prevalence of the *H*_*i*_ strategy if all players were only participating in the *k*-th hawk dove game and other parts of their macro strategy did not contribute to fitness. In order to represent macro strategies, we use a binary notation. Macro strategy *j* corresponds to a binary number *t*_*j*_, where the *i*-th element of *t*_*j*_ (*t*_*j,i*_) is equal to 1(0) if it involves playing hawk (dove) in game *i*. For example, if macro strategy *j* is *H*_1_ *D*_2_ *D*_3_ *H*_4_ then *t*_*j*_ = (1, 0, 0, 1).

We define the pay–off of playing the hawk strategy in game *k* at time *t* as *R*_*k,H*_(*t*) (*R*_*k,D*_(*t*) for playing dove). Furthermore, 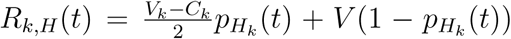 (and the analogous equation for the dove). The time derivative of *s*_*H*_*j*__is obtained from the standard two-player hawk dove replicator equation and is equal to 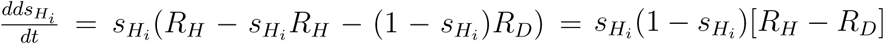. Based on the definition of our model, we know that the pay-off of the macro strategy j is given by *R*_*j*_(*t*) =∑*i t*_*j,i*_*R*_*k,H*_ (*t*) + (1 − *t*_*j,i*_)*R*_*k,D*_(*t*).

With the above notation, we write our guess for solution to the overall replicator equation as

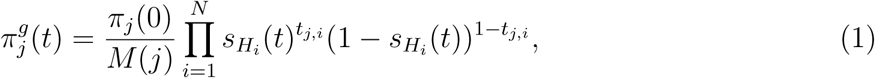

with *M* (*j*) a normalization factor for strategy *j*, chosen so that at time *t* = 0, the above equation gives the initial condition value *π*_*j*_(0). This trial solution can be examined by taking a time derivate of Equation (1), giving

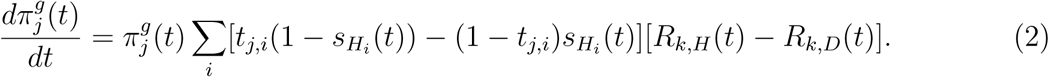

On the other hand, the replicator equation states that 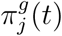 must satisfy

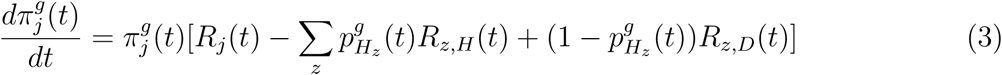

If we expand out the terms for *R*_*j*_(*t*) and replace 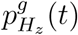 with 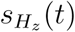 (we justify this below), then we can see that Equations (3) and (2) are equivalent and therefore (1) is the solution to our model.

The key to this calculation is the assumption that 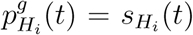. To show why this can many terms, namely the *π*_*j*_(0), the *M* (*j*) and all the *s*_*H*_*i*__. Note that since all the variables were approximately be the case, we write: 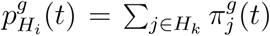. Each term 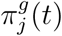 is a product of initialized independently, we can think of this as a product of independent random variables and we can calculate the expected value of the product by multiplying together the expected values of all terms. When we carry out this procedure, we find exactly that *p*_*H*_*i*__(*t*) = *s*_*H*_*i*__(*t*) in expectation.

We verified with computational simulations that Equation (1) produces very good approximations to our model. In Figure S1, you can see that the simulated equilibrium probabilities of our model match very well the approximate analytical solution.

**Figure 3:**
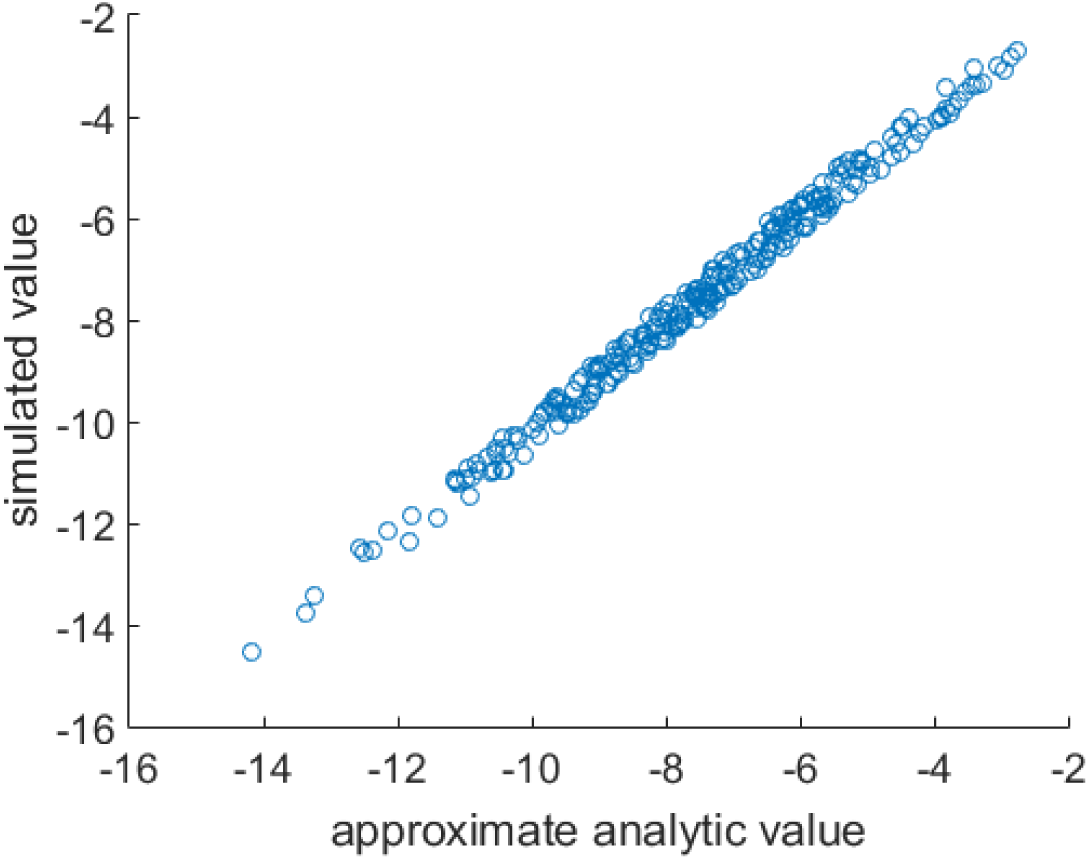
The analytical formula in Equation (1) gives a good approximation to the simulated results. Probabilities obtained from the approximate analytic Equation (1) and equilibrium solution in simulation. This is a log-log plot to facilitate seeing the entire range of probabilities.

### Log-normality

The approximate analytical solution provides a key insight into the model. First, note that Equation (1) may be thought of as a product of multiple random variables. While the sum of many random variables tends to a Gaussian, the product of many random variables tends to a log-normal distribution. This helps explain why we observe log-normal distributions in our simulations.

### Multi-modality

As mentioned in the main text, certain SADs may be weakly multi-modal. Using our analytical approximation, we can see that one way in which multi-modality can emerge is if one of the *V*_*i*_ values is much larger than all the others (close to C as opposed to close to 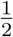). Because the *V*_*i*_ interact multiplicatively, this causes the product probabilities to split into two groups: one group is containing the term 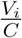 in it, which is approximately 1, the other group contains the term 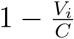, which is close to zero. As you can see in Figure 4, multi-modal distributions can thus emerge in the SAD. Biologically, this might correspond to a competitive scenario where one type of resource has very low value for survival.

**Figure 4:**
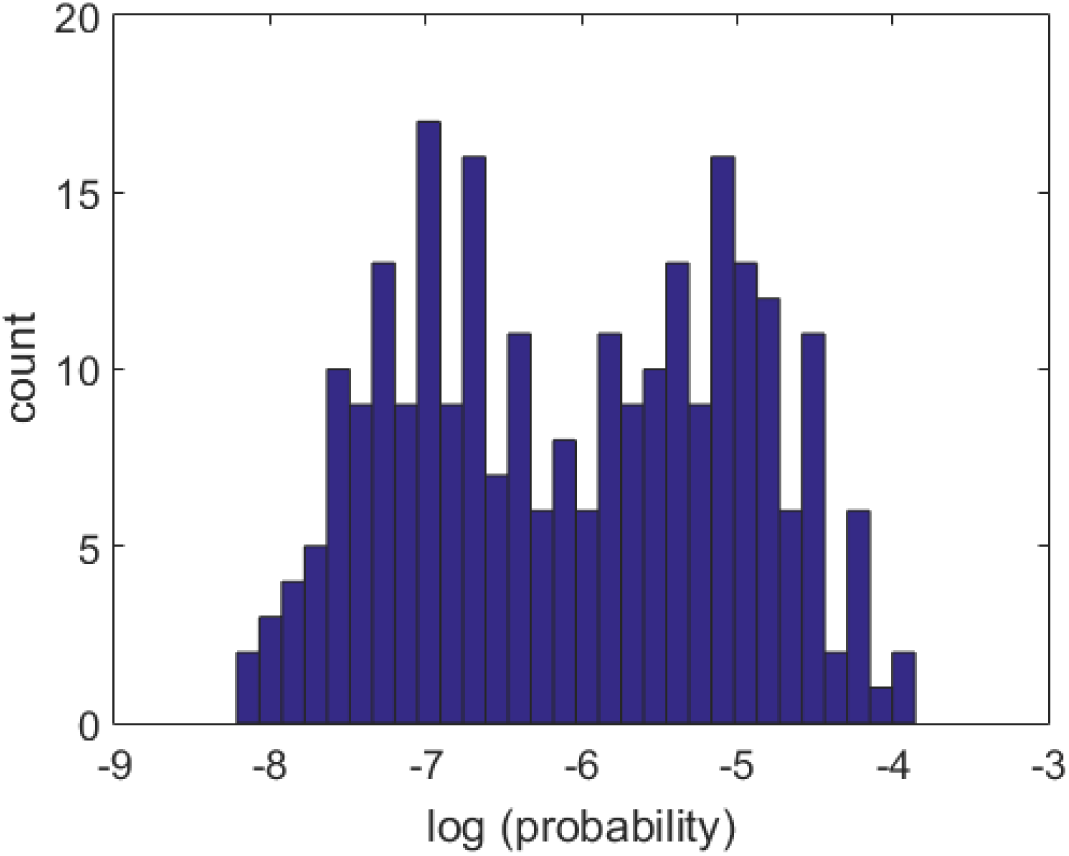
Example histogram of a simulation which converged to a multi-modal distribution. For this simulation, *C* = 2, *V*_1_ = 1.9 and *V*_2_ to *V*_8_ was sampled uniformly at random between 1 and 1.1.

